# Systematic characterization of zinc in a series of breast cancer cell lines reveals significant changes in zinc homeostasis

**DOI:** 10.1101/2025.01.11.632547

**Authors:** Mena Woyciehowsky, Portia Larson, Annika R. Stephan, Sharee L. Dandridge, Doreen Idonije, Kylie A. Berg, Alyx Lanthier, Stephanie Araiza Acuna, Saskia W. Stites, Waverly J. Gebhardt, Samuel E. Holtzen, Ananya Rakshit, Amy E. Palmer

## Abstract

An optimal amount of labile zinc (Zn^2+^) is essential for proliferation of human cells, where Zn^2+^ levels that are too high or too low cause cell cycle exit. Tumors of the breast have been characterized by high levels of total Zn^2+^. Given the role of Zn^2+^ in proliferation of human cells and elevation of zinc in breast cancer tumors, we examined the concentration of total and labile Zn^2+^ across a panel of 5 breast cancer cell lines, compared to the normal MCF10A cell line. We found that three cell lines (MDA-MB-231, MDA-MB-157, and SK-Br-3) showed elevated labile Zn^2+^ in the cytosol, while T-47D showed significantly lower Zn^2+^, and MCF7 showed no change compared to MCF10A cells. There was no change in total Zn^2+^ across the cell lines, as measured by ICP-MS, but we did observe a difference in the cells ability to accumulate Zn^2+^ when Zn^2+^ in the media was elevated. Therefore, we examined how proliferation of each cell line was affected by increases and decreases in the media. We found striking differences, where three cancer cell lines (MDA-MB-231, MDA-MB-157, and MCF7) showed robust proliferation in high Zn^2+^ at concentrations that killed MCF10A, T-47D, and SK-Br-3 cells. We also discovered that 4 of the 5 cancer cell lines demonstrate compromised proliferation and increased cell death in low Zn^2+^, suggesting these cells may be addicted to Zn^2+^. Overall, our study suggests significant differences in Zn^2+^ homeostasis and regulation in different types of breast cancer cells, with consequences for both proliferation and cell viability.

## Introduction

Zinc (Zn^2+^) is an essential micronutrient that is necessary for proliferation of human cells; ^1^ decreased Zn^2+^ abrogates proliferation, while subtle increases in Zn^2+^ increase proliferation. However, high concentrations of Zn^2+^ can compromise proliferation in human cells. A recent study showed that there is a pulse of labile Zn^2+^ in human cells during the cell cycle, where Zn^2+^ increases from ∼200 pM to 1.6 nM for about 2 hrs in early G1.^2^ If this pulse is too small (in Zn^2+^ depleted conditions) or too large (in Zn^2+^ excess conditions), cells pause the cell cycle until the optimal Zn^2+^ concentration is re-established. These studies reveal that Zn^2+^ plays a crucial role in regulating the cell cycle, and hence proliferation of human cells. Moreover, these studies show that the amount of Zn^2+^ present in growth media affects proliferation.

In addition to proliferation, Zn^2+^ plays a role in differentiation, metabolism, DNA synthesis, transcriptional regulation, chromatin organization, and genome stability.^3–5^ At the molecular level, Zn^2+^ is a cofactor in thousands of proteins, including all six major enzyme classes and 48% of human transcription factors.^6^ Cellular concentrations of Zn^2+^ are controlled by a network of transporters, transcription factors, and the buffering protein, metallothionein. The average amount of total Zn^2+^ in a mammalian cell is in the hundreds of micromolar,^7^ but most of it is bound to proteins such that the labile or exchangeable pool is in the hundreds of picomolar range.^8–10^ However, the labile pool is dynamic and can change as a function of cellular processes,^2^ signaling events,^4,11–13^ or in disease states.^9,14–17^

Numerous human cancers are characterized by changes in serum and tumor zinc levels. Decreased zinc has been documented in the serum of patients with cancer of the breast, gallbladder, colon, bronchus, or lung.^18^ Tumors have been characterized by both decreases in zinc (prostate cancer) and increases in zinc (breast and lung cancer).^18,19^ Moreover, human cancers have also been characterized to have altered levels of Zn^2+^ transporters.^19–24^ In many of these cases, transporter expression and/or Zn^2+^ levels track with tumor aggressiveness, suggesting that altered Zn^2+^ homeostasis may contribute to the severity of cancer.^20,22^

The role of zinc in breast cancer has been particularly enigmatic. While there have been conflicting reports in the literature, a meta-analysis of 36 studies on zinc in breast cancer found consensus that serum and plasma levels were significantly lower in breast cancer patients compared to healthy controls, and zinc in cancerous breast tissue was found to be significantly higher.^25^ These studies typically involve measuring the total zinc in a tissue sample by elemental analysis techniques such as laser-ablation inductively coupled mass spectrometry (LA-ICPMS), or x-ray fluorescence microscopy. Tumor samples are heterogeneous, consisting of cancer cells, stroma, extracellular matrix, and perhaps infiltrated immune cells,^26^ making it difficult to determine what part of the tumor exhibits elevated Zn^2+^. There have been a handful of studies that have attempted to measure the labile Zn^2+^ pool in breast cancer cell lines using fluorescent indicators. However, these studies have relied on small molecule sensors which have been shown to have heterogeneous localization.^9,27^ Moreover, few of these studies carried out an *in situ* calibration,^28^ which is required to make conclusions about relative Zn^2+^ levels. The one exception is a recent study that systematically compared three different small molecule sensors (TSQ, FluoZin-3, and ZinPyr-1) in a series of breast cancer cell lines. Importantly, the authors carried out calibrations and determined that ZinPyr-1 was most appropriate for quantifying labile Zn^2+^ and determined the labile Zn^2+^ pool to be higher in breast cancer cell lines than the non-cancerous MCF10A cell line (MDA-MB-231 (1.25 nM) > MCF7 (1.02 nM) >T-47D (0.74 nM) > MCF10A (0.62 nM)).^27^ However, as the authors noted, ZinPyr-1 localized to the cytosol as well as intracellular organelles and the localization appeared different in different cell lines, with some showing stronger perinuclear fluorescence than others. Therefore, the authors cautioned that the labile Zn^2+^ pool estimates were likely a combination of Zn^2+^ in the cytosol and organelle pools. To our knowledge, no study has used a genetically encoded fluorescent sensor that can be rigorously targeted to the cytosol in breast cancer cell lines to define the cytosolic Zn^2+^ pool explicitly.

Given the role of Zn^2+^ in cell proliferation,^1,29^ genome stability,^3,5,30^ the documentation of altered Zn^2+^ homeostasis in human breast cancer, and the well-known aberrant proliferation of cancer cells, we set out to examine Zn^2+^ in a series of breast cancer cell lines. We selected cancer cell lines with different molecular characteristics (basal versus ductal, activation of different hormone receptors) and compared them to non-cancerous MCF10A cells. We created stable cell lines expressing a genetically encoded fluorescent sensor to measure labile Zn^2+^ in the cytosol and discovered that many breast cancer cells have higher labile Zn^2+^ than MCF10A cells with the following order MDA-MB-231 ∼ MDA-MB-157 > SK-Br-3 > MCF7 ∼ MCF10A > T-47D. However, we detected no difference in the total Zn^2+^ levels by ICP-MS when cells were grown in minimal media. When media was supplemented with ZnCl_2_, there were profound differences in Zn^2+^ accumulation, suggesting differences in Zn^2+^ homeostasis. We also found that cancer cells responded differently to Zn^2+^ in the media with respect to proliferation. Some cancer cell lines appeared to thrive in high Zn^2+^ at concentrations that killed MCF10A cells. In addition, a number of cancer cell lines appeared to be addicted to Zn^2+^ such that mild Zn^2+^ deficiency induced cell death. Combined, our results reveal that zinc homeostasis is significantly altered in breast cancer cells but different types of breast cancer cells have very different phenotypes.

## Results

Breast cancer is heterogeneous disease.^31^ At the molecular level, breast cancers are typically characterized by proliferation status (luminal A vs luminal B), activation status of the human epidermal growth factor receptor 2 (HER2), estrogen receptor (ER), and progesterone receptor (PR), as well as mutational status of *BRCA1* and *BRCA2*.^31–33^ There are multiple connections between zinc and these proteins. ER, PR, and BRCA1 are zinc-dependent transcription factors. Estrogen and progesterone have been shown to alter the expression of zinc transporters, leading to changes in zinc.^34,35^ Zinc has been shown to influence kinase signaling cascades downstream of HER2.^23,36^ But whether different molecular characteristics affect cellular Zn^2+^ or Zn^2+^ homeostasis is not clear. One study found higher Zn in ER+ tumors compared to ER-tumors by X-ray Fluorescence Microscopy.^37^ However, this was directly contradicted by a study which used laser ablation ICP-MS to measure total zinc in a series of tumors, which showed that triple negative tumors (ER- PR-HER2-) showed the highest levels of Zn, with HER2+ tumors also showing high Zn.^38^ For this study we selected 5 cancer cell lines that span a range of molecular characteristics, as defined by activation of hormone receptors (**Figure 1A**). All cell lines have wild type BRCA1.^39^

**Figure 1:**
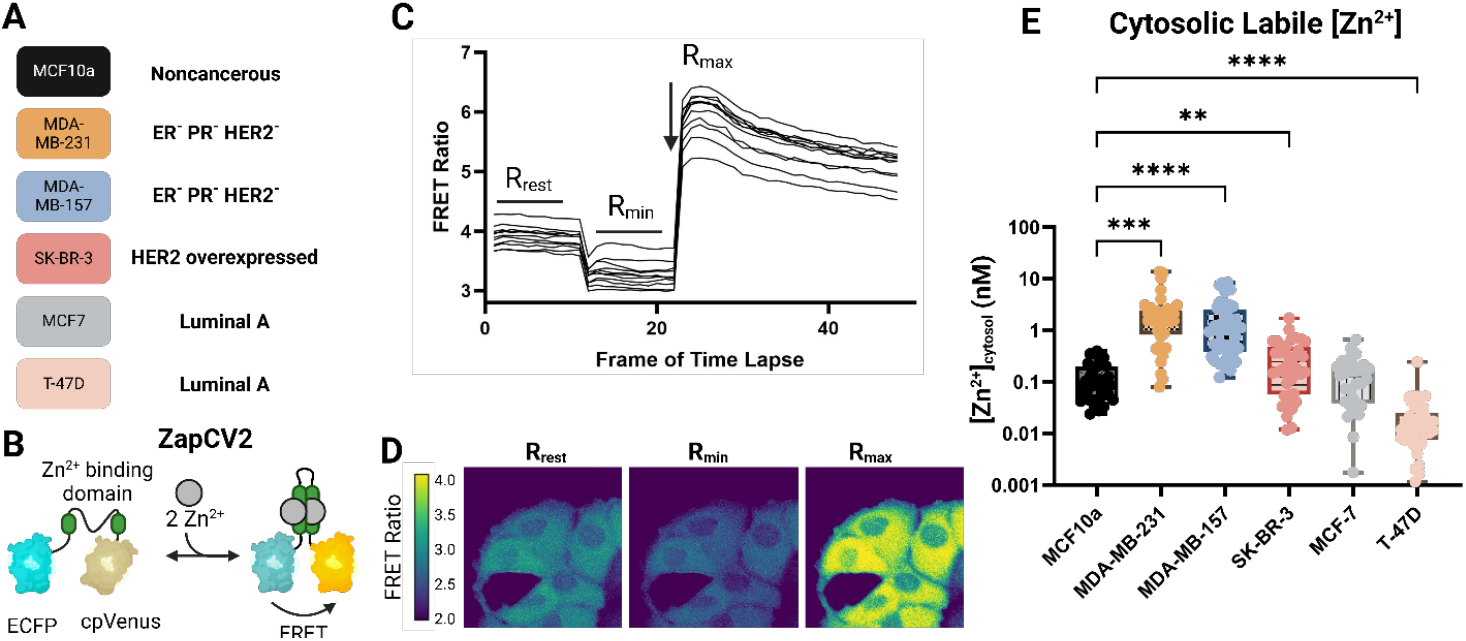
Quantification of cytosolic Zn^2+^ in selected breast cancer cell lines using the genetically encoded FRET sensor ZapCV2. A. Selected breast cancer cell lines included in this study and their respective molecular subtypes. Each cell line was stably transfected with both NES-ZapCV2 and H2B-HaloTag. B. Schematic of the ZapCV2 sensor illustrating that Zn^2+^ binding increases FRET from ECFP to cpVenus. The ratio of FRET to CFP is proportional to the Zn^2+^ concentration. C. A representative ZapCV2 calibration in MCF7 cells. Cells are imaged in HEPES-buffered Hank’s balanced salt solution (HHBSS), then treated with R_min_ solution containing the Zn^2+^ chelator TPA to acquire the FRET ratio of the fully desaturated sensor. The chelator is then washed out and an R_max_ solution containing buffered Zn^2+^ and pyrithione is added to acquire the FRET ratio of the fully saturated sensor. D. Representative images of each phase of the calibration in MCF7 cells. R_rest_ corresponds to the FRET ratio of cells in HHBSS with no treatment. R_min_ corresponds to cells treated with 50 µM TPA. R_max_ corresponds to cells treated with buffered Zn^2+^ and pyrithione. Scale bar corresponds to 20 µm. E. The calculated cytosolic Zn^2+^ for each cell line in nM. Statistical significance was determined *via* Brown-Forsythe and Welch ANOVA test with Dunnett’s T3 comparison test (^**^*p* < 0.01; ^***^*p* < 0.001; ^****^*p* < 0.0001) for n ≥ 3 biological replicates.

To quantify the labile Zn^2+^ pool in the cytosol, we used PiggyBac transposase to create cell lines stably expressing the genetically encoded Zn^2+^ sensor NES-ZapCV2 (**Figure 1B-C**).^1,40^ This sensor is composed of a Zn^2+^ binding domain sandwiched between enhanced cyan fluorescent protein (ECFP) and circularly permuted Venus. When two Zn^2+^ ions bind to the sensor, a conformational change shifts the pair of fluorescent proteins closer, increasing the efficiency of Förster Resonance Energy Transfer (FRET). Cells were subjected to an *in situ* calibration to determine the minimum and maximum FRET ratio in each cell and these parameters were used to determine the concentration of labile Zn^2+^ in the cytosol in each cell (**Figure 1D, Supplementary Table 1**). **Supplementary Figure 1** reports the fractional saturation and dynamic range of the sensor in each cell line. Three of the five cancer cell lines had higher cytosolic Zn^2+^ than MCF10A cells, with the two triple negative cell lines having the highest levels (2.5 nM and 1.8 nM for MDA-MB-231 and MDA-MB-157, respectively). The next highest was the HER2+ cell line SK-Br-3 (310 pM). The cytosolic Zn^2+^ in MCF7 cells was comparable to MCF10A cells (150 pM and 124 pM, respectively). Finally, the T-47D breast cancer cell line had significantly lower Zn^2+^ than MCF10A cells (24 pM vs 124 pM). These results reveal that labile Zn^2+^ is indeed different in cancer cell lines, but different cell lines may show higher or lower Zn^2+^ compared to MCF10A cells. The highest Zn^2+^ levels were found in triple negative and HER2+ cell lines.

We next examined total Zn using ICP-MS. When cells were grown in minimal media we found no difference in the total Zn reported as fg/cell (**Figure 2A, 2B, Supplementary Table 2**). However, when cells were grown in media supplemented with different amounts of zinc, significant differences emerged. Addition of 30 μM ZnCl_2_ to the media for 48 hr prior to isolation of cells for ICP-MS led to a 2.2-fold increase in the average total Zn in MCF10A cells but did not lead to a substantial increase in any of the breast cancer cell lines, suggesting differences in their ability to regulate Zn^2+^ homeostasis. Treatment of cells with 150 μM ZnCl_2_ for 48 hrs led to an increase in average total Zn in MCF7 and SK-Br-3 cells (6.4-fold increase and 7.7-fold increase, respectively), but no change in the total Zn level in T-47D, or MDA-MB-157 cells. MDA-MB-231 cells showed a small, but statistically significant 1.3-fold increase in average total Zn in 150 μM ZnCl_2_. The total Zn in MCF10A cells could not be measured because treatment with 150 μM ZnCl_2_ led to cell death. These results suggest that the cells respond differently to zinc in their environment, with some cell lines resisting accumulation of excess zinc.

**Figure 2:**
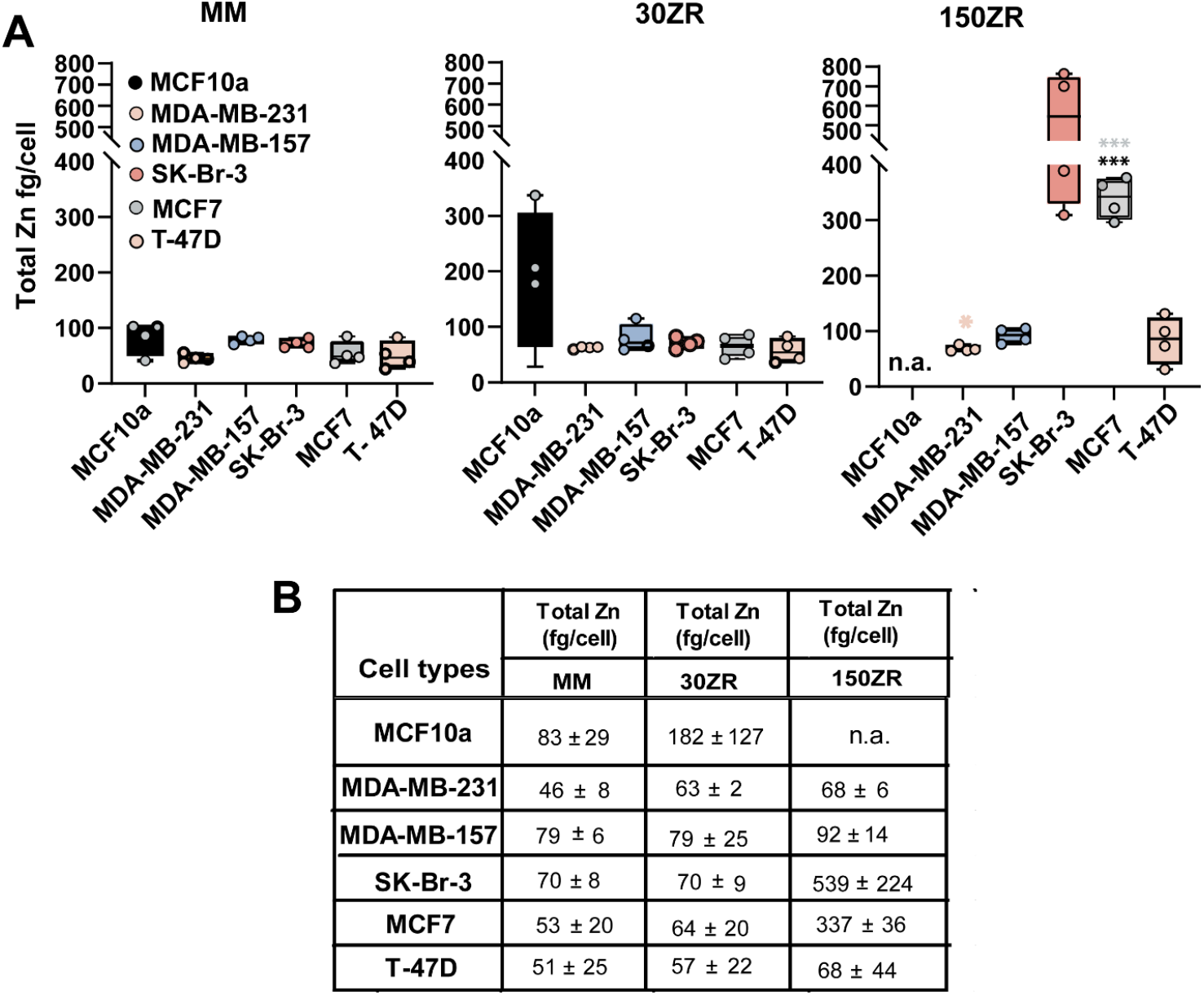
Breast cancer cells have variable amount of total zinc in different zinc media. (A) ICP-MS analysis of total zinc pool in non-cancerous MCF10a cells and different breast cancer cells grown for 48h in minimal media (MM), zinc rich (ZR) conditions 30ZR (MM + 30 µM ZnCl_2_) and 150ZR (MM + 150 µM ZnCl_2_). All measurements were performed in four biological replicates (N=4). Error bars represent SEM. Significance was determined *via* ANOVA with Tukey’s multiple comparison test (^***^*p* < 0.001; ^****^*p* < 0.0001). Black and colored asterisks represent statistical significance of differences with respect to MCF10a MM condition and MM conditions of each cell types respectively. MCF10a cells were all dead under 150ZR conditions after 48h. (B) Table reflects the average values of total zinc pool(fg/cell) in each cell types.

Cellular zinc homeostasis is intricately interconnected with the regulation of other biologically relevant metals. To explore these connections, we used ICP-MS to investigate both basal levels as well as changes in the total amount of Cu, Fe, Mn, Ca, Co, Mg, P, and S (**Supplementary Figure 2, Table 3 & 4)**. ICP-MS analysis revealed that MCF10A and T-47D cells grown in minimal media (MM) had the highest total Cu levels (4.2 fg/cell and 6.7 fg/cell, respectively), while MDA-MB-231 cells exhibited the lowest (1.4 fg/cell) **(Supplementary Figure 2A)**. In addition, zinc supplementation (30ZR) increased total Cu levels in MCF10A and MCF7 (1.2-fold and 2.5-fold, respectively) cells. Total Cu levels remain elevated in MCF7 cells under 150ZR conditions. However, total Cu in other cancer cells remained unchanged indicating differential regulation of Cu metabolism in response to zinc availability. Total Fe levels in MCF10A (27.1 fg/cell), MDA-MB-157 (26.6 fg/cell), and MCF7 (29.1 fg/cell) were comparable, whereas other cancer cells showed lower levels with MDA-MB-231 showing the lowest (9.4 fg/cell). MDA-MB-231 cells have 2.8 & 2.9-fold lower iron than MDA-MB-157 and MCF10A respectively (**Supplementary Fig 2B, Table 3)**. However, zinc supplementation had no significant impact on total Fe levels. MCF10A cells had the lowest total Mn levels under MM (0.6 ± 0.2 fg/cell) whereas MDA-MB-231 and MDA-MB-157 cells had significant 4-fold and 6.6-fold higher levels of total Mn compared to MCF10A cells (**Supplementary Figure 2C, Supplementary Table 3 & 4)**. T-47D, SK-Br-3, and MCF7 cells had similar levels of total manganese in MM. Surprisingly, zinc supplementation consistently decreased total Mn levels across all cell lines, with reductions of up to 2.2-5.3-fold in MCF7, SK-Br-3, and T-47D cells under 150ZR, suggesting an inverse relationship between zinc and Mn accumulation. MCF10A showed at least 3-fold higher levels of Co than MCF7 and MDA-MB-231 but 2.2-fold lower levels than SK-Br-3 cells (**Supplementary Figure 2E, Supplementary Table 3)**.

We also looked at the total levels of non-trace elements including calcium (Ca), magnesium (Mg), sulfur (S), and phosphorous (P) levels reported in pg/cell in breast cancer cells and compared them to MCF10A cells following zinc supplementation. We noted that total Ca and Mg levels were highest in MDA-MB-157 cells in MM (0.27 pg/cell and 1.05 pg/cell, respectively), while MCF10A exhibited the lowest levels (0.02 pg/cell and 0.46 pg/cell) (**Supplementary Figure 2D & 2F, Supplementary Table 3 & 4)**. SK-Br-3, MCF7, and MDA-MB-231 cells showed comparable total Ca and Mg levels. Zinc supplementation had no significant effect on Ca or Mg levels across cell lines. No significant changes were observed in total sulfur and phosphorus levels across cell lines (**Supplementary Figure 2G and 2H)**. Overall, these findings highlight distinct baseline profiles of elemental homeostasis across breast cancer cell lines and demonstrate that zinc supplementation differentially alters total Cu and Mn levels while having minimal effects on total Fe, Ca, Mg, and other metals.

Because cells responded differently to zinc in their environment, we examined whether the cell lines proliferated differently in response to zinc in the media. To measure cell proliferation, we plated the same number of cells in each well of a 96-well plate, subjected different wells to minimal media containing the Zn^2+^ chelator tris(2-pyridylmethyl)amine (TPA) or increasing amounts of ZnCl_2_ for 48 hrs. We then used a resazurin assay to measure the conversion of resazurin to fluorescent resorufrin in each well, where a higher signal corresponded to more cells with metabolically active mitochondria. **Figure 3A** shows that MCF10A cells have decreased proliferation in zinc-deficient media (2 μM TPA, 2ZD; and 3 μM TPA, 3ZD), a slight increase in proliferation in a moderate zinc rich media (30 μM ZnCl_2_, 30ZR), and compromised proliferation in rich zinc media (100 μM ZnCl_2_, 100ZR and above). This matches previous observations.^1,2^ **Figure 3B** and **C** show a similar decrease in proliferation for the triple-negative breast cancer cell lines MDA-MB-231 and MDA-MB-157 in zinc-deficient media. However, in high zinc, these cells either continue to proliferate comparable to minimal media (MDA-MB-231) or show only a slight decrease at very high zinc (250 ZR and above, MDA-MB-157). **Figure 3D** shows the SK-Br-3 cell line exhibits decreased proliferation in zinc-deficient media, and progressive compromise of proliferation in elevated zinc, starting at 150ZR. The MCF7 cells behave similarly to the triple-negative breast cancer cell lines, showing a decrease in proliferation in zinc-deficient media, and no perturbation of proliferation in high zinc until 500ZR (**Figure 3E**). Finally, as shown in **Figure 3F** the T-47D cells don’t tolerate perturbation from minimal media; they show compromised proliferation in zinc-deficient and zinc-rich media.

**Figure 3:**
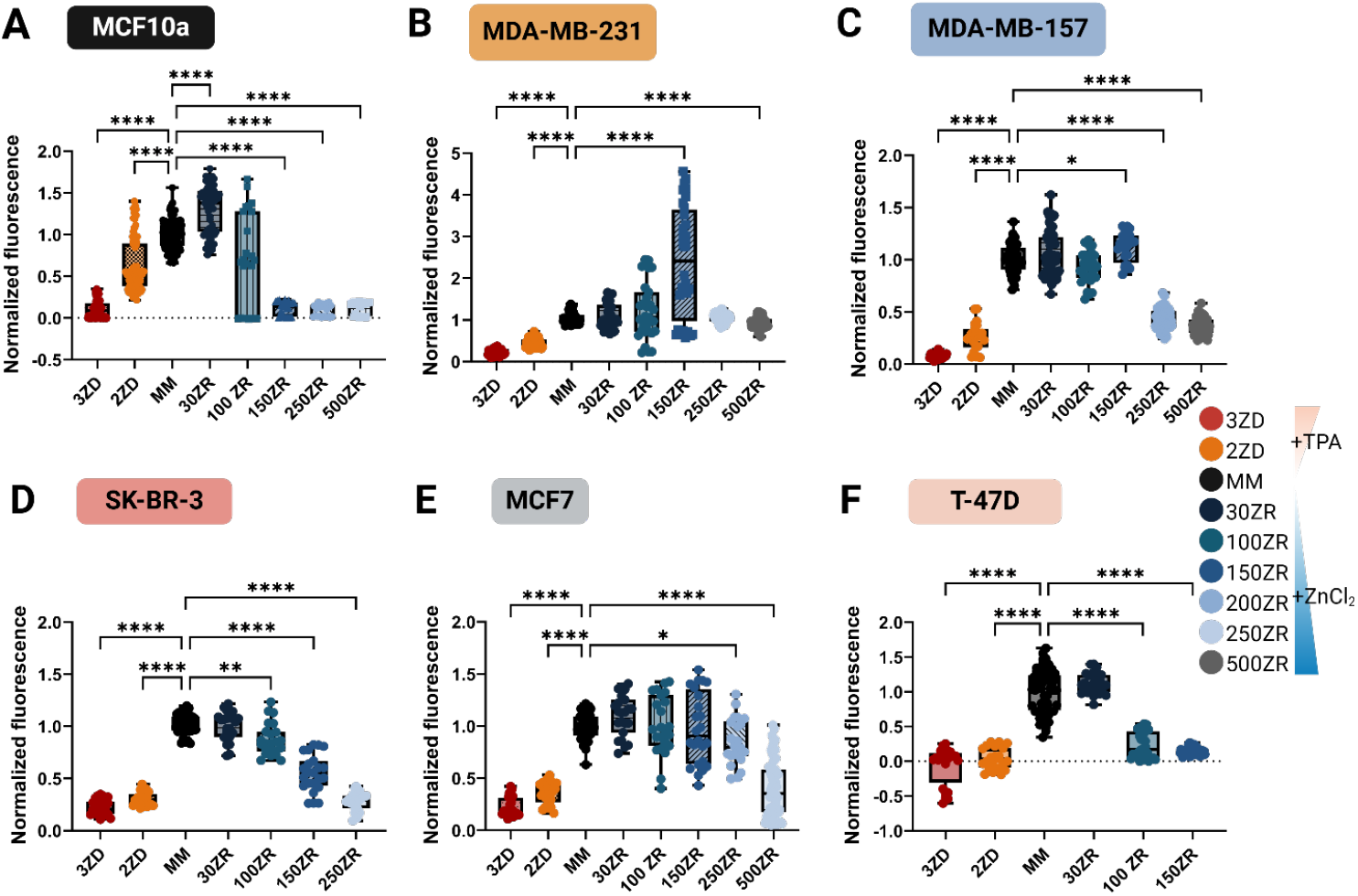
Proliferation of breast cancer cells changes in response to the available Zn^2+^ in minimal media. Results of resazurin cell proliferation assay on MCF10A (A), MDA-MB-231 (B), MDA-MB-157 (C), SK-BR-3 (D), MCF7 (E), and T-47D (F) cells grown for 48 hours in minimal media supplemented with varying levels of TPA and ZnCl_2_. Each data point represents one well of cells. The fluorescence counts in each well are normalized to the average fluorescence counts in the minimal media condition and statistical significance was determined with respect to the minimal media (MM) condition for each cell type. 3ZD = 3 μM TPA, 2ZD = 2 μM TPA, 30ZR = 30 μM ZnCl_2_, 100ZR = 100 μM ZnCl_2_, 150ZR = 150 μM ZnCl_2_, 250ZR = 250 μM ZnCl_2_, 500ZR = 500 μM ZnCl_2_. Statistical significance was determined *via* Brown-Forsythe and Welch ANOVA with Dunnett’s T3 multiple comparison test (\**p* < 0.05; ^**^*p* < 0.01; ^****^*p* < 0.0001) for n ≥ 3 biological replicates.

While the resazurin assay provides information on how the density of cells changes over time in different growth conditions, it can’t differentiate between a situation in which cells go quiescent and stop dividing from a situation in which cells die off in each growth condition. Therefore, we carried out a ReadyProbes Cell Viability assay. Cells were grown for 48 hrs in minimal media with either 2 μM TPA, 30 μM ZnCl_2,_ or 150 μM ZnCl_2_ and stained with NucBlue and NucGreen to identify the total number of cells and number of dead cells, respectively. **Figure 4A** shows that MCF10A cells have compromised viability in high Zn (150ZR) but were largely unaffected in 30ZR or 2ZD. This reveals that the low proliferation in **Figure 3A** in 2ZD is not the result of cell death, but rather the cells exiting the cell cycle and going quiescent, as shown previously.^1,2^ **Figure 4B** and **C** show that the two triple-negative breast cancer cell lines thrive in high Zn^2+^ as there is no increase in cell death in the 30ZR or 150ZR conditions. However, these cells are extremely sensitive to subtle decreases in Zn^2+^, as 2ZD caused a significant increase in the fraction of dead cells. This result suggests that the MDA-MB-231 and MDA-MB-157 don’t exit the cell cycle in low Zn^2+^ like MCF10A cells. **Figure 4D** shows that SK-Br-3 cells show a slight increase in cell death in 150ZR, but no increase in cell death in low Zn^2+^ (2ZD), suggesting these cells behave similar to MCF10A cells. **Figure 4E** shows that MCF7 cells exhibit increased cell death in 2ZD and 150ZR, suggesting that compromised proliferation is due to increased cell death rather than cell cycle exit. Finally, in **Figure 4F**, T-47D shows high cell death in both low and high Zn^2+^, consistent with proliferation data suggesting this cell line can’t tolerate changes in Zn^2+^ in the media.

**Figure 4:**
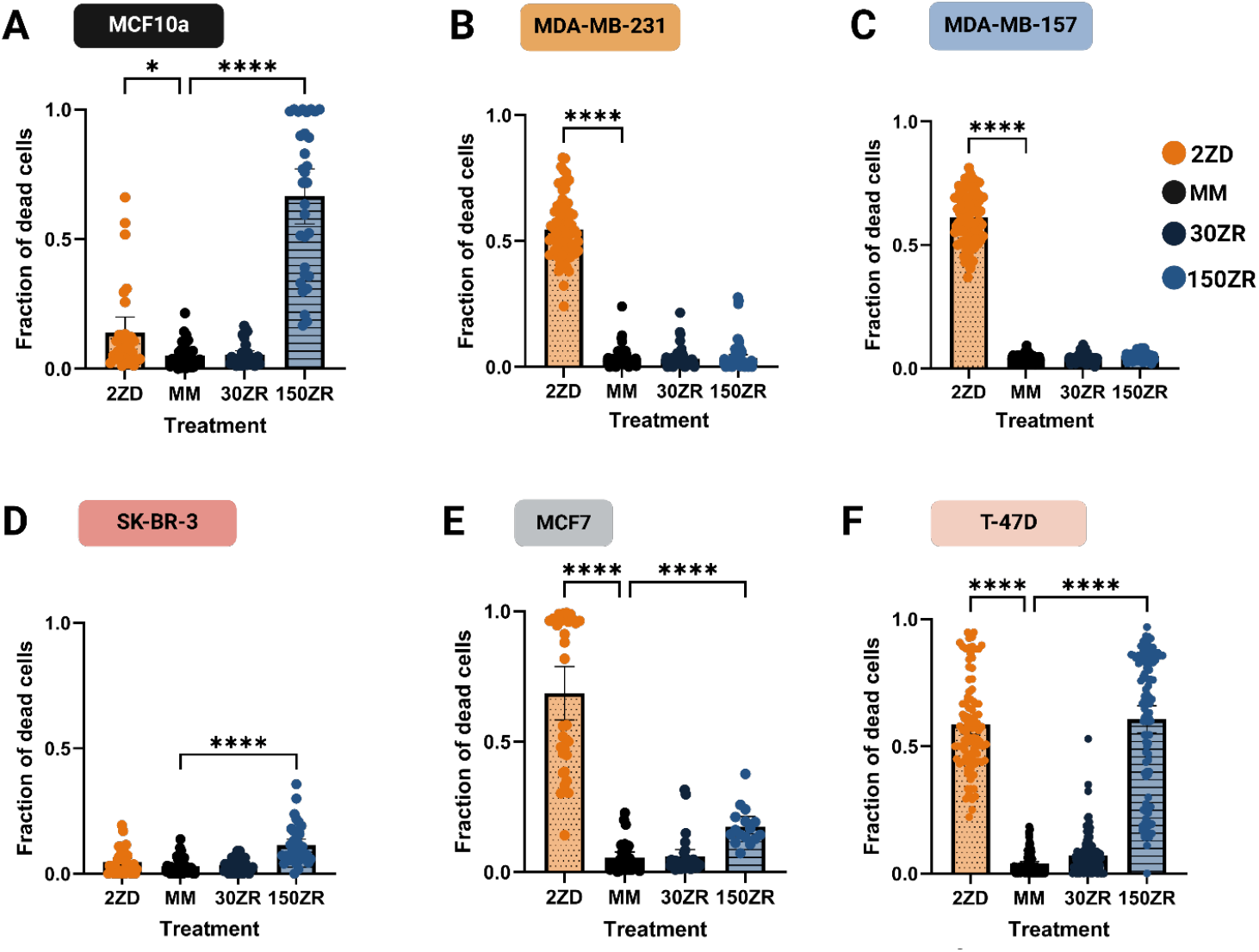
Viability of breast cancer cells changes with available Zn^2+^ in media. Results of Live/Dead assay on MCF10a(A), MDA-MB-231(B), MDA-MB-157(C), SK-BR-3(D), MCF7 (E), and T-47D(F) cells grown for 48 hours in media containing 1.5% chelex-100 treated serum (minimal media, MM) supplemented with 2 µM TPA (2ZD), 30 µM ZnCl_2_ (30ZR), and 150 µM ZnCl_2_ (150ZR). All measurements were performed in two biological replicates (N = 2) and statistical analysis were performed *via* Brown-Forsythe and Welch ANOVA with Dunnett’s T3 multiple comparison test (Significance: ^*^*p* < 0.05, ^***^*p* < 0.001; ^****^*p* < 0.0001). Error bars represent SEM.

The proliferation assay in **Figure 3** was carried out in a defined minimal media with limited serum so that we could rigorously change the zinc availability. We did control experiments to verify that the minimal media did not significantly alter proliferation compared to full-growth media (**Supplementary Figure 3**). Still, we wanted to define whether cells responded differently to zinc perturbations in full-growth media containing 10% serum compared to minimal media. We hypothesized that perhaps they would be less sensitive to ZD and ZR conditions because the serum would help buffer Zn^2+^ availability. **Figure 5** shows the results of the resazurin proliferation assay after growth for 48 hours in full-grown media (FGM) with different amounts of TPA or ZnCl_2_. MCF10A cells still showed compromised proliferation under conditions of zinc deficiency (2ZD and 3ZD) but showed more robust proliferation in zinc-rich conditions, up until 200ZR, above which cells stopped proliferating (**Figure 5A**). The triple-negative breast cancer cells (MDA-MB-231 in **Figure 5B** and MDA-MB-157 in **Figure 5C** showed compromised proliferation in zinc deficiency, but slightly increased proliferation in zinc-rich conditions, even up to 500ZR. Again, these results suggest profoundly altered zinc homeostasis in these cell lines. The HER2+ SK-Br-3 cell line showed compromised proliferation in zinc-deficient conditions, and stable proliferation in zinc-rich conditions until 500ZR which showed a marked decrease in proliferation (**Figure 5D**). The MCF7 cell line showed no change in proliferation in low Zn^2+^ and very little change in proliferation in high Zn^2+^, with a slight decrease starting at 200ZR (**Figure 5E**). Finally, the T-47D cell line showed no change in proliferation until 500ZR which showed a slight decrease in proliferation.

**Figure 5:**
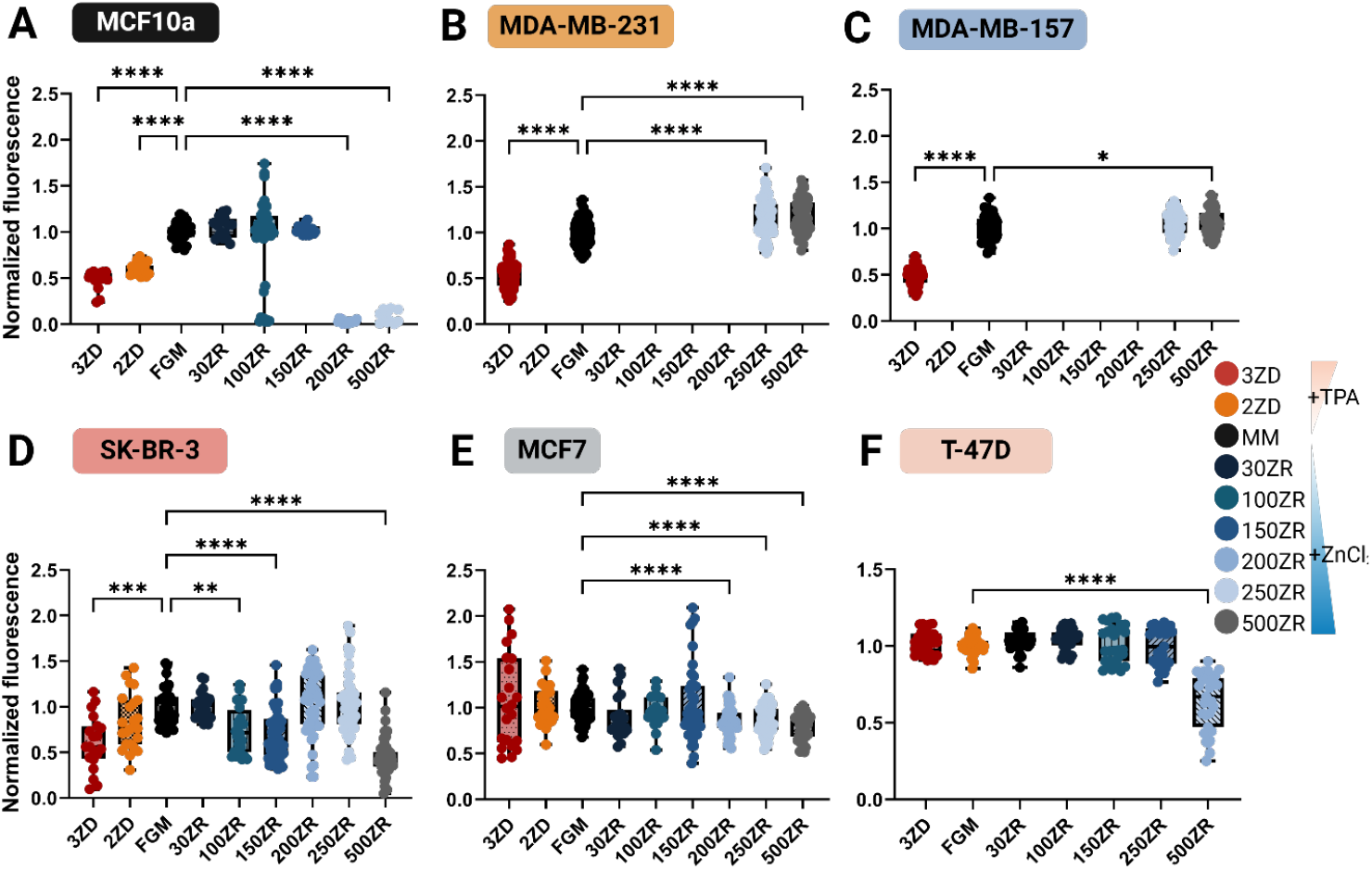
Proliferation of breast cancer cells changes in response to available Zn^2+^ in full growth media. Results of resazurin cell viability assay on MCF10A (A), MDA-MB-231 (B), MDA-MB-157 (C), SK-BR-3 (D), MCF7 (E), and T-47D (F) cells grown for 48 hours in full growth media supplemented with varying levels of TPA and ZnCl_2_. Each data point represents one well of cells. The fluorescence counts in each well are normalized to the average fluorescence counts in the full growth media condition. Statistical analysis was performed via Brown-Forsythe and Welch ANOVA with Dunnett’s T3 multiple comparison test (Significance^****^ = p < 0.001, ^***^ = p < 0.001, ^**^ = p < 0.01, ^*^ = p < 0.05. n ≥ 3 biological replicates)

## Discussion

Zinc plays a critical role in cell cycle regulation and proliferation making zinc homeostasis essential for proper execution of Zn-dependent biological processes in all human cell sub-types.^1,2,5^ Previous studies have shown that labile Zn^2+^ is dynamic and fluctuations in labile Zn^2+^ levels are integral to cell cycle progression and hence proliferation.^2^ However, the relationship between Zn^2+^ homeostasis and proliferation of breast cancer cells was unclear due to conflicting findings on total and labile Zn^2+^ levels in breast cancer cells and tissues.^25^ In this study, we explored the role of Zn^2+^ in the proliferation of luminal, HER2++, and triple-negative breast cancer cells by systematically altering Zn^2+^ availability in the growth media. We used a combination of genetically encoded fluorescent sensors, ICP-MS, and cell proliferation assays to examine labile Zn^2+^ pools, total Zn, and the effects of changes in zinc supplementation across breast cancer cell lines with heterogeneous HER2, ER, and PR expression levels. Our study highlights distinct Zn^2+^ homeostasis profiles across breast cancer cell lines and non-cancerous MCF10A cells, emphasizing the importance of labile Zn^2+^ levels, metal homeostasis, and cellular proliferation in cancer cell physiology.

There is consensus across multiple studies that breast cancer tissue contains elevated levels of zinc. But given the heterogeneity of cancer tissue, this does not necessarily mean that breast cancer cells contain elevated zinc. Moreover, most prior studies measured total, rather than labile zinc. The labile Zn^2+^ pool is of particular interest because this pool of zinc is unbound and hence biochemically available to influence cell physiology. In this study we quantified total and labile zinc across a series of different breast cancer cell lines. Surprisingly, in minimal media we found no changes in total zinc, as measured by ICP-MS, suggesting that elevated zinc in breast cancer tissue may come from changes in zinc in the extracellular matrix, stromal cells, infiltrated immune cells, or other. However, we did find differences in the labile Zn^2+^ pool. Specifically, three cancer cell lines (MDA-MB-231, MDA-MB-157, and SK-Br-3) showed elevated labile Zn^2+^ in the cytosol compared to MCF10A and one cell type (T-47D) showed significantly lower Zn^2+^. Finally, MCF7 showed no change in labile Zn^2+^ compared to MCF10A. The high levels of Zn^2+^ in MDA-MB-231 compared to MCF10A is consistent with a previous which used a small molecule fluorescent Zn^2+^ indicator (ZinPyr-1) to measure Zn^2+^ in MDA-MB-231, MCF7, T-47D, and MCF10A cells.^27^ However, the relative amount of Zn^2+^ and the concentration of Zn^2+^ in MCF7 and T-47D cells differs in our study compared to this previous study. A notable difference in our approach is that in this study we used a genetically encoded sensor explicitly targeted to the cytosol of cells, so our concentrations report on the concentration of Zn^2+^ in the cytosol. ZinPyr-1 shows heterogeneous localization across the four cell types, with some fluorescence in the cytosol, as

well as fluorescence in a perinuclear region that appears very similar to the Golgi apparatus (in MCF10A and MDA-MB-231 cells) or in vesicles (in MCF7 and T-47D cells).^27^ Therefore, the differences could arise from changes in Zn^2+^ in organelles.

Given that breast cancer tumors show increased levels of total zinc, but cancer cells, at least those measured in this study, don’t have elevated total zinc, we speculated that perhaps in tumors cancer cells experience a high zinc environment due to increased zinc in extracellular matrix or surrounding cells. Therefore, we examined how changes in zinc availability in growth media affect proliferation of cells. Previously, we have shown that in zinc deficient conditions MCF10A cells exit the cell cycle and go quiescent. However, they don’t die. Rather, when Zn^2+^ is resupplied, the cells re-enter the cell cycle in a synchronous fashion. Here we found that all cancer cell lines show compromised proliferation in low Zn^2+^, as measured by lower cell densities in a resazurin assay. Decreased cell densities could result from cell cycle exit, like MCF10A cells or could be due to cell death. We found that 4 of the 5 cancer cell lines (MDA-MB-231, MDA-MB-157, MCF7, and T-47D) show increased cell death in low Zn^2+^ (**Figure 3, 4**). The final cancer cell line SK-Br-3 showed compromised proliferation but no increase in cell death, suggesting that similar to MCF10A, these cells exit the cell cycle and go quiescent in low Zn^2+^. The increased cell death in low Zn^2+^ suggests that the cancer cell lines are addicted to Zn^2+^.

There are also significant differences in how cancer cells manage elevated Zn^2+^ compared to each other and compared to MCF10A cells. When MCF10A cells are treated with 30 μM ZnCl_2_ in the media, they show a slight increase in proliferation. It is important to note that increases in Zn^2+^ in the media can lead to increases in the cytosol, but limited Zn^2+^ transport across the plasma membrane and other homeostasis mechanism buffer how much the intracellular labile Zn^2+^ pool increases. For example, we have shown previously that 30 μM ZnCl_2_ in the media leads to an increase in cytosolic labile Zn^2+^ from ∼ 100 pM to 75 nM in MCF10A cells.^30^ Increases in Zn^2+^ above 30 μM lead to compromised proliferation (**Figure 3A**) and increased cell death (**Figure 4A**) for MCF10A cells. 4 of the 5 cancer cell lines are more tolerant of elevated Zn^2+^, with the 3 of the 5 (MDA-MB-231, MDA-MB-157, and MCF7) appearing to thrive in very high Zn^2+^ (up to 250 μM ZnCl_2_ in the media), conditions that kill MCF10A cells. SK-Br-3 and T47-D are more sensitive to high Zn^2+^ and show increased cell death at 150 mM ZnCl_2_ in the media, similar to MCF10A cells. Overall, our results suggest that some breast cancer cell lines (MDA-MB-231, MDA-MB-157, and MCF7) have significantly altered Zn^2+^ homeostasis compared to MCF10A, SK-Br-3, and T-47D; they can’t tolerate low Zn^2+^, which leads to cell death and they thrive in high Zn^2+^, under conditions that kill MCF10A, and many other cell types.

The changes in labile Zn^2+^ and response to Zn^2+^ in the media suggest changes in proteins that regulate zinc homeostasis, namely Zn^2+^ importers (slc39a1-14, ZIP1-14), Zn^2+^ exporters (slc30a1-10, ZnT1-10), and buffering proteins (metallothioneins). To gain insight into how zinc regulatory proteins might be altered in the different cell lines, we turned to a published study which performed quantitative proteomics on the cancer cell line encyclopedia, which includes MDA-MB-231, MDA-MB-157, MCF7, and T-47D.^41^ While there are 14 Zn^2+^ importers, only 8 were detected in the cancer cell lines (ZIP1, ZIP4, ZIP6, ZIP7, ZIP8, ZIP10, ZIP11, and ZIP14). Similarly, while there are 10 Zn^2+^ exporters, only 5 were detected (ZnT1, ZnT5, ZnT6, ZnT7, and Znt9). Across all the transporters, that largest difference between the breast cancer cells was an increase in expression of ZnT1 in MDA-MB-231 cells compare to the average across all cancer cell lines, and a decrease in expression in T-47D cells. This could explain how MDA-MB-231 cells tolerate high Zn^2+^, they simply export it back into the media, and why T-47D are intolerant of high Zn^2+^, with downregulated ZnT1, they are incapable of exporting elevated Zn^2+^. The study also showed higher levels of metallothionein expression in MDA-MB-231 cells (MT1L, MT1G, and MT2A) compared to the average across all cancer cell lines, and a decrease in metallothionein in T-47D. Again, this would be consistent with T-47D cells being unable to buffer increases in the labile Zn^2+^ pool, making them more susceptible to Zn^2+^ toxicity. Overall, all the breast cancer cells show a decrease in Zip4 and an increase in Zip7 compared to the average expression across all cell lines.

In summary, we report a systemic study of several breast cancer cell subtypes using genetically encoded sensors, ICP-MS, and biochemical assays to define labile Zn^2+^, total Zn^2+^ and other trace metals, proliferation, and cell viability conditions where Zn^2+^ availability in the media is altered. Our results suggest significant changes in Zn^2+^ regulation and homeostasis, with consequences for cell proliferation and cell viability. We found that cancer cell lines with different molecular phenotypes (luminal vs basal, triple negative) had significantly different Zn^2+^ phenotypes.

## Methods

### Cell culture

MCF10a, MDA-MB-231, MDA-MB-157, MCF7, SK-Br-3 and T-47D cells were acquired from ATCC and maintained in full growth medium (FGM). MCF10a cells were grown in FGM containing DMEM/F12 medium supplemented with 5% horse serum, 1% pen/strep antibiotics, 20 ng/mL EGF, 0.5 μg/mL hydrocortisone, 100 ng/mL cholera toxin, and 10 μg/mL insulin at 37

°C, 5% CO_2_ and 90% humidity. MCF-7 cells were maintained in FGM consisting of RPMI 1640 media supplemented with 10% fetal bovine serum (FBS) and 1% penicillin/streptomycin at 37 °C and 5% CO_2_. SK-BR-3 cells were maintained in FGM consisting of McCoys5a L-Glutamine media supplemented with 10% FBS and 1% penicillin/streptomycin at 37 °C and 5% CO_2_. Both MDA-MB-231 and MDA-MB-157 cells were maintained in Leibovitz’s L15 media supplemented with 10% fetal bovine serum (FBS), and 1% Pen/strep. These two cell lines were maintained in an incubator at 37 °C with 0% CO_2_. T-47D cells were grown in FGM containing RPMI 1640, 10% FBS and 1% penicillin/streptomycin at 37 °C, 5% CO_2_ and 90% humidity. All cells except MDA-MB cells were passaged with 0.05% trypsin-EDTA solution. MDA-MB cells were passaged with 0.25% trypsin-EDTA solution. Imaging and ICP-MS experiments, and some proliferation experiments, used minimal medium (MM) which was designed to sustain normal proliferation under controlled amount of zinc in the media. MM usually consists of Chelex-100 treated serum. MM for all breast cancer cell line contains 1.5% chelex treated FBS, 1% Pen/Strep along with the main component of the media. MM for MDA-MB cells: L15 media, chelexed 1.5% FBS, and 1% pen/strep; MM for MCF7: RPMI 1640 media with glutamax, 1.5% chelexed FBS, and 1% pen/strep; MM for SK-Br-3: McCoys5a media, 1.5% chelexed FBS, and 1% pen/strep; MM for T-47D: RPMI 1640 media 1.5% chelexed FBS 1% pen/strep and 0.02% chelexed insulin. MM for MCF10a consists of 50:50 Ham’s F12 phenol red free/FluoroBrite DMEM with 1.5% Chelex 100-treated horse serum, 10 μg/mL Chelex 100-treated insulin, 1% pen/strep antibiotics, 20 ng/mL EGF, 0.5 μg/mL hydrocortisone, and 100 ng/mL cholera toxin. Proliferation and viability assays were performed in zinc deficient (ZD), zinc rich(ZR), minimal media(MM) and FGM solution. Zinc-rich (ZR) and zinc-deficient (ZD) media conditions were prepared by adding the following to MM: 15 µM ZnCl_2_ (15ZR), 30 µM ZnCl_2_ (30ZR), 50 µM ZnCl_2_ (50 ZR), 100 µM ZnCl_2_ (100ZR) 150 µM ZnCl_2_ (150ZR), or 250 µM ZnCl_2_ (250ZR), 500 µM ZnCl_2_ (500ZR), 2 µM TPA (2ZD) or 3 µM TPA (3ZD). All cells were plated in FGM and after 24 hours, the media was changed to zinc conditions listed above. After 48h, cells were subjected to the resazurin assay to determine cell proliferation in the different media conditions.

### Generation of stable cell lines

Stable cell lines were generated by electroporating plasmids into the cells. The MCF10A cell line was generated previously in Palmer laboratory.^1^ Breast cancer cells MDA-MB-231, MDA-MB-157, SK-Br-3, MCF7 and T-47D were electroporated with super-PiggyBac transposase (SPBT, XXX), PB-NES-ZapCV2, and PB-H2B-HaloTag plasmids to generate stable cell lines expressing the Zn^2+^ specific sensor NES-ZapCV2 and nuclear marker H2B-HaloTag. PB-NES-ZapCV2 and PB-H2B-HaloTag plasmids were inserted at different times to ensure proper expression of each plasmid in every single cell. Wildtype breast cancer cells were grown in respective media at 80-100% confluency and trypsinized before electroporating using Neon electroporator. Each electroporation reaction on 1 × 10^6^ –2 × 10^6^ cells required 500 ng of SPBT, and either 2000 ng of PB-NES-ZapCV2 or 2000 ng of PB-H2B-HaloTag plasmids. Electroporation on each breast cancer cell line was performed following Neon Protocols from ThermoFisher Scientific. Cells expressing PB-NES-ZapCV2 and PB-H2B-HaloTag plasmid were selected with Blasticidin and G418 antibiotics, respectively. Successful expression of NES-ZapCV2 was confirmed by visualizing fluorescence either in the CFP or YFP channel. H2B-HaloTag expression was confirmed by incubating cells with JF646-HaloTag ligand which binds to the histone 2B protein resulting in nuclear fluorescence. Stable cells expressing both plasmids showed fluorescence corresponding to both NES-ZapCV2 and H2B-HaloTag. These stable breast cancer cells were then used for experiments discussed in this paper.

### Measurement of labile Zn^2+^ using the genetically encoded NES-ZapCV2 sensor

To measure the labile Zn^2+^ concentration in cancer cells and MCF10A cells, we conducted *in-situ* sensor calibration in cells expressing NES-ZapCV2 sensor.^28^ Zinc calibration involves a sequential measurement of the resting FRET ratio (R_rest_) of the NES-ZapCV2 sensor, followed by measurement of the minimum FRET ratio (the lower limit of ZapCV2 sensor, R_min_) upon addition of a cell permeable zinc chelator, TPA. This was then followed by measurement of the maximum FRET ratio (the upper limit of ZapCV2 sensor, R_max_) by adding Zn^2+^, saponin, and pyrithione to the cells. About 200,000-500,000 cells were plated in MM in a 35 mm imaging dish and grown for 24 hours before *in situ* calibration. Cells were washed three times with PO_4_ ^3-^free HEPES-buffered Hanks’ Balanced Salt Solution (HHBSS) and equilibrated at 25 ^o^C for 30 mins before the experiment. For FRET ratio measurements, cells were imaged on a Nikon-HCA microscope and fluorescence was measured in the CFP-YFP FRET and CFP channels, keeping similar laser power and exposure time. Parameters for Zn^2+^ calibration experiments were optimized for each cell type. To measure R_rest_, FRET and CFP intensity were measured at 30 s intervals for 5-8 minutes. Then, 100 μM TPA in PO_4_^3-^ free HHBSS was added to the cells in PO_4_ ^3-^ free HHBSS to a final concentration of 50 μM TPA, resulting in the measurement of R_min_ at 30 s intervals for 5 mins. Cells were then washed twice with PO_4_ ^3-^, Ca^2+^, and Mg^2+^-free HHBSS (pH 7.4) solution to remove excess TPA. Cells were imaged again for 5-8 mins at 10 s intervals after the addition of 59.6 nM buffered Zn^2+^ solution, 0.001% saponin, and 750 nM pyrithione in PO_4_ ^3-^ free HHBSS to obtain the maximum FRET ratio, R_max._ Filter sets (excitation = Ex, emission = Em) used were as follows: CFP Ex: 440, 455 dichroic, Em: 480/20, power 30-50, exposure 200-400 ms; YFP Ex: 508, 518 dichroic, Em: 540/21; CFP/YFP FRET Ex: 440, 455 dichroic, Em: 540/21, power 30-50, exposure 200-400ms. All experiments for each cell type were performed in replicates (n ≥ 3). The data were analyzed using ImageJ and MATLAB programs. R_rest_ and R_min_ correspond to the average FRET ratio in resting and TPA treated conditions respectively, whereas R_max_ represents the maximum FRET ratio. The FRET ratio was calculated using following equation:

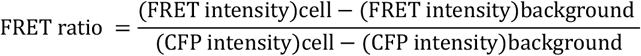

After getting R_rest_, R_min_ and R_max_, free zinc was calculated by the following equation:

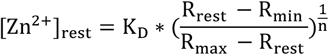

where K_d_, dissociation constant for ZapCV2 was 5.3 ± 1.1 nM and Hill coefficient n was 0.29<u>^10^</u> and dynamic ratio (DR) of the sensor was calculated as

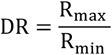

Fractional saturation (FR) of the sensor was calculated as

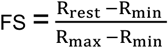

For calculating labile Zn^2+^, cells with a fractional saturation (FR) between 0 and 1, and dynamic range (DR) >1.35 were included in the analysis. Statistical significance was determined *via via* Brown-Forsythe and Welch ANOVA test with Dunnett’s T3 comparison test (\**p* < 0.05; ^**^*p* < 0.01; ^***^*p* < 0.001; ^****^*p* < 0.0001).

### Elemental analysis of total metals via ICP-MS

ICP-MS was performed to measure the total Mg, Ca, P, S, Mn, Fe, Co, Cu, and Zn amount in cells grown in defined MM with and without Chelex-100 treated FBS, HS, and insulin. Cells were grown in either MM, 30ZR and 150ZR media for 48h. Cells were spun down and washed twice with 1X PBS prepared in Chelex-100 treated water. Cells were then counted four times before digestion. Cell pellets containing 0.06-9 million cells were vortexed for 3 mins in 100 µL of elemental grade TraceSELECT Ultra nitric acid and digested for 2h at 90 ^o^C on a heat-bath. After digestion, the mixture was diluted with 900 µL of 1% HNO_3_ in chelexed dd H_2_O. All ICP-MS measurements were performed in the OHSU Elemental Analysis Core and results were normalized to average cell counts.

### Resazurin assay

The proliferation of MCF10a, MDA-MB-231, MDA-MB-157, MCF7, SK-Br-3 and T-47D cells grown under different zinc conditions was measured using Resazurin assay. This assay was performed on cells grown in different zinc conditions, prepared either in minimal media (MM) or full growth media (FGM). MCF10A and breast cancer cells were plated in 96 well plates at a cell density of 5000-10,000 cells/well in FGM. For the resazurin assay in MM, media was changed to defined zinc media after 24h and then cells were grown for 48h. For measurements in FGM, the media was replaced with defined zinc media after 24h and cells were incubated in zinc supplemented FGM for 48h. After 48h, 20 μL of 0.55 mg/mL aqueous resazurin dye was added to the media in each well bringing the final volume to 220 μL and cells were incubated for 4h at 37 ^o^C. After incubation, fluorescence from the 96 well plate was measured using a plate reader (λ_ex_: 550 ± 5 nm, λ_em_: 590 ± 5 nm, integration time: 300 ms). For each biological replicate, cells in each zinc condition were plated in 8 technical replicates. The fluorescence intensity of each condition was first background subtracted and outliers were determined using an Interquartile Range (IQR) test. After removing outliers, corrected data from zinc treated conditions were then normalized to the average value from either MM or FGM control conditions. Statistical significance was determined by *via* Brown-Forsythe and Welch ANOVA test with Dunnett’s T3 comparison test (\**p* < 0.05; ^**^*p* < 0.01; ^***^*p* < 0.001; ^****^*p* < 0.0001).

### Live/Dead assay

Cell viability in MCF10A cells and breast cancer cells was investigated using Invitrogen ReadyProbes^TM^ Cell Viability Imaging Kit (Blue/green). Cells were plated in 96 wells plates at 5000/well cell density in full growth media (FGM) and media was changed to MM, 2ZD, 30ZR and 150ZR after 24h. Cells were then grown in different zinc supplemented minimal media for 48h. Cells were then stained with 28 µL mixture of blue/green staining solution for 20 mins and imaged on Nikon HCA microscope using DAPI and GFP channel. Cells in each zinc conditions were plated in ≥ 6 technical replicates for each biological replicate and every experiment resulted in imaging at 3 three regions of interest (ROI). The imaged cells were counted using in-house codes in MATLAB. The ratio of dead cells to the total number of cells was used as a metric of cell morbidity. All measurements were performed in two biological replicates (N = 2) and statistical analysis was performed *via* Brown-Forsythe and Welch ANOVA test with Dunnett’s T3 comparison test.

## Author contributions

M.W and P.L contributed equally to this work. M.W, P.L., S.D., A.R., and A.E.P conceptualized and developed methodologies. A.R., and A.E.P wrote and edited the manuscript with inputs from all authors. M.W, P.L., S.D., and K.B developed stable cell lines. M.W, P.L, S.D., S.E.H, and W.G performed zinc calibration assays. M.W., P.L., S.D., K.B., D.I., and S.A.A designed and carried out resazurin assays. A.L and S.S performed live-dead assays. M.W, P.L, and A.R performed ICP-MS experiments and analyzed the data. S.E.H wrote all the MATLAB codes necessary to analyze calibration data and live-dead assay. Data analysis and figures: M.W., P.L., A.S., S.D., K.B., D.I., S.E.H., A.R and A.E.P. Funding acquisition, supervision, and project administration, A.E.P.

## Acknowledgements

We thank the University of Colorado Biochemistry Cell Culture Core Facility specifically Theresa Nahreini, for providing resources for cell culturing. We would like to thank the OHSU Elemental Analysis Core facility especially Sophia Miller and Dr. Martina Ralle for performing ICP-MS experiments. ICP-MS measurements were performed in the OHSU Elemental Analysis Core with partial support from NIH (S10RR025512). We thank Dr. Martina Ralle for useful discussion on ICP-MS analysis. We would like to acknowledge NIGMS MIRA R35 GM139644 (A.E.P), the Biological Sciences Initiative (BSI, CU Boulder), and Undergraduate Research Opportunity Program (UROP, CU Boulder) for generous financial support.

## Notes

### Competing Interest Statement

The authors have declared no competing interest.

