## Supplementary information for "Systematic characterization of zinc in a series of breast cancer cell lines reveals significant changes in zinc homeostasis"

**Supplementary information contents:**

Supplementary Figure 1: Fractional saturation and dynamic range of the NES-ZapCV2 sensor in each breast cancer cell line.

Supplementary Figure 2: ICP-MS analysis of trace elements in non-cancerous MCF10a and different breast cancer cell types.

Supplementary Figure 3: Resazurin assay for cell proliferation under different media conditions.

Supplementary Table 1: Quantification of labile cytosolic $Zn^{2+}$

Supplementary Table 2: Statistical significance of the differences in total zinc from Brown-Forsythe and Welch ANOVA with Dunnett's T3 multiple comparison test between MCF10a and different breast cancer cells in different zinc conditions via ICP-MS analysis.

Supplementary Table 3: Total amount of non-zinc trace elements in non-cancerous MCF10A cells and different breast cancer cells in different zinc conditions via ICP-MS analysis.

Supplementary Table 4:  Statistical significance of the differences in total non-zinc trace elements between MCF10a and different breast cancer cells in different zinc conditions via ICP-MS analysis.

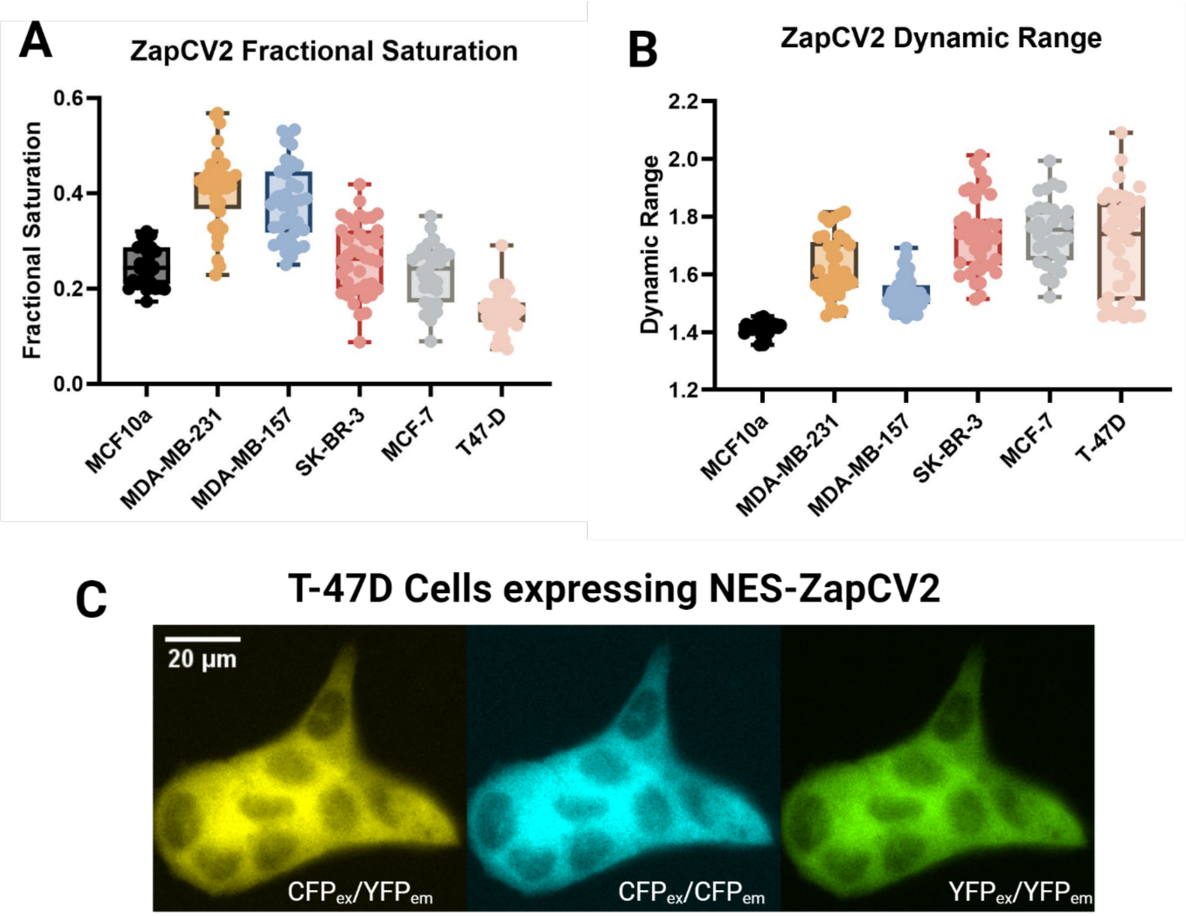

**Supplementary Figure 1: Fractional saturation and dynamic range of the NES-ZapCV2 sensor in each breast cancer cell line.** A. The fractional saturation of the NES-ZapCV2 sensor at rest for each breast cancer cell line and noncancerous control. B. The dynamic range of the NES-ZapCV2 sensor in each breast cancer cell line and noncancerous control. C. Representative images of nuclear excluded (NES) ZapCV2 expressing T-47D cells. Images represent FRET (yellow), CFP (cyan) and YFP (green) in the same field of view. Scale bar corresponds to 20 µm.

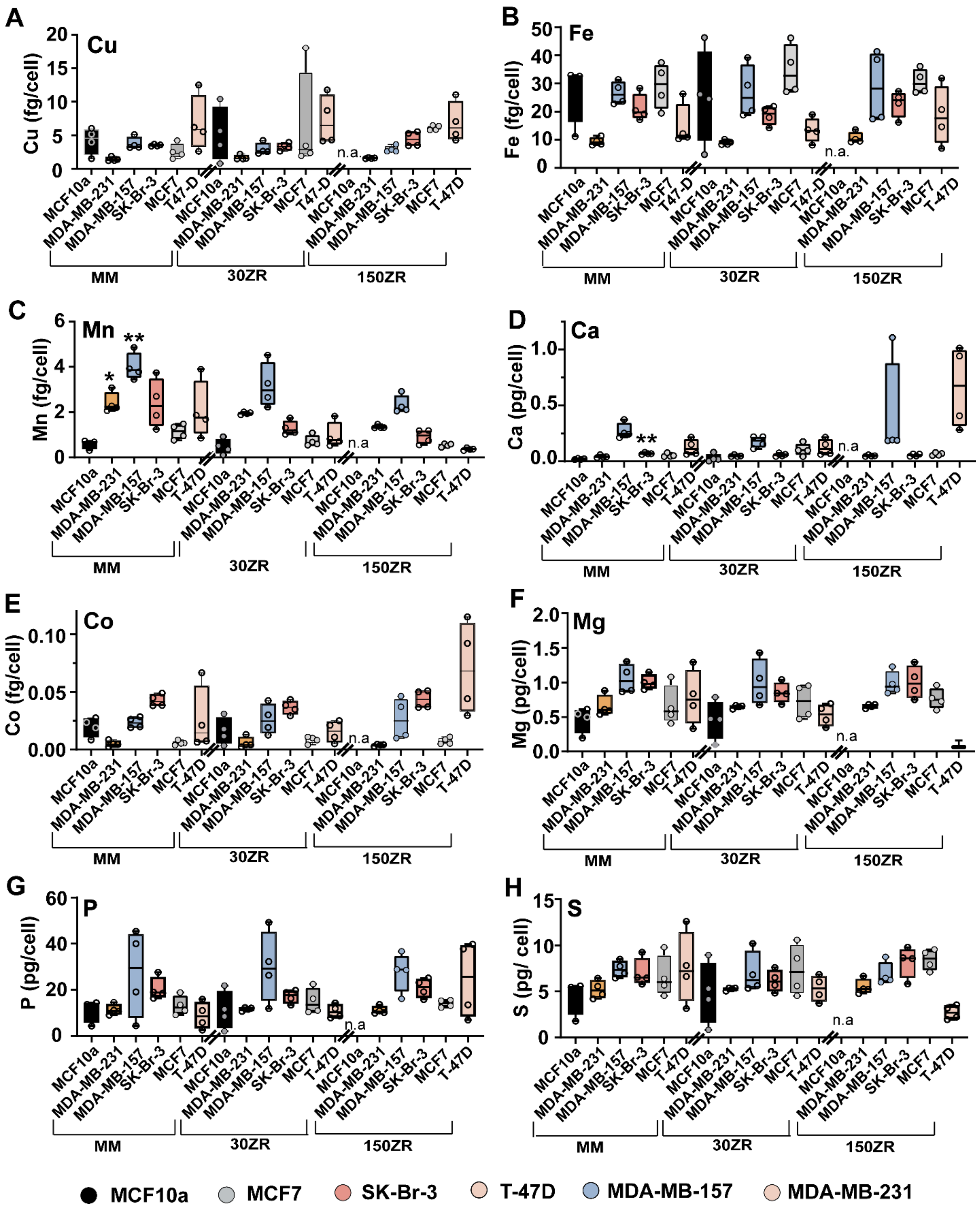

**Supplementary Figure 2: ICP-MS analysis of trace elements in non-cancerous MCF10a and different breast cancer cell types**. (A-H) Measurements of total amount of copper (Cu), iron (Fe), manganese (Mn), calcium (Ca), cobalt (Co), magnesium (Mg), sulphur (S) and phosphorous

(P) in cells grown in Minimal media (MM), 30ZR (MM + 30 μM $ZnCl_2$) and 150ZR (MM + 150 μM $ZnCl_2$) conditions for 48h. All measurements were performed in four biological replicates (N = 4) and statistical analysis were performed. Error bars represent SEM. Significance was determined *via* Brown-Forsythe and Welch ANOVA with Dunnett's T3 multiple comparison test ($^{*}p < 0.05$; $^{**}p < 0.01$; $^{***}p < 0.001$; $^{****}p < 0.0001$). MCF10A cells were all dead in 150ZR conditions after 48h. Significance of results for minimal media condition of different cell types were calculated with respect to MCF10A MM condition.

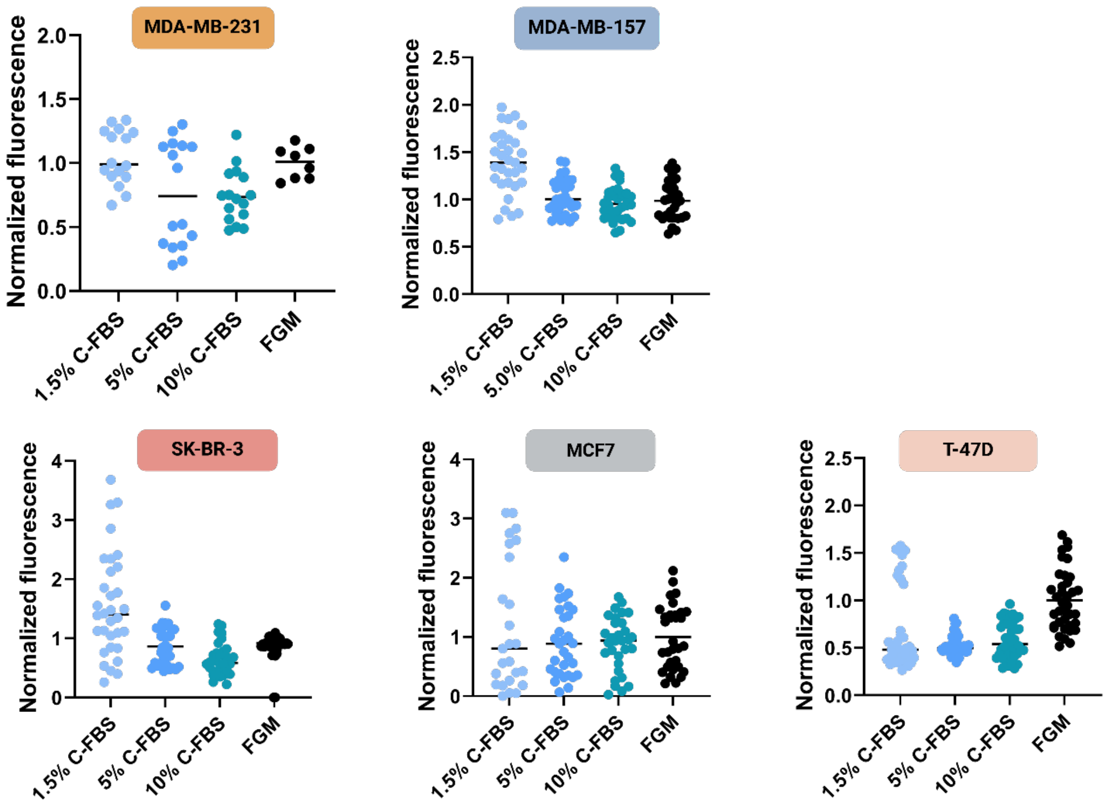

**Supplementary Figure 3: Resazurin assay for cell proliferation under different media conditions.** Cells were grown for 48 hrs in the following media: full growth media containing 10% FBS (FGM), growth media plus 10% Chelex-treated FBS (10% C-FBS), growth media plus 5% Chelex-treated FBS (5% C-FBS), or growth media plus 1.5% Chelex-treated FBS (1.5% C-FBS). Each dot represents one well in a 96 well plate. The fluorescence intensity in each well is normalized to the average fluorescence intensity of the full growth media condition. There were no significant differences in proliferation in FGM versus media will Chelex-treated serum across any of the breast cancer cell lines.

**Supplementary Table 1: Quantification of labile cytosolic $Zn^{2+}$**

| | MCF10a | MDA-MB-231 | MDA-MB-157 | SK-BR-3 | MCF7 | T47D |
|---|---|---|---|---|---|---|
| Number of values | 35 | 37 | 43 | 40 | 31 | 41 |
| Minimum | 0.02378 | 0.08001 | 0.1209 | 0.01181 | 0.001744 | 0.001161 |
| 25% Percentile | 0.04498 | 0.8113 | 0.3805 | 0.05645 | 0.03838 | 0.007509 |
| Median | 0.08595 | 1.65 | 1.024 | 0.1793 | 0.1312 | 0.01517 |
| 75% Percentile | 0.1964 | 2.445 | 2.503 | 0.4747 | 0.1934 | 0.02517 |
| Maximum | 0.397 | 13.65 | 8.469 | 1.706 | 0.6503 | 0.2439 |
| Range | 0.3732 | 13.57 | 8.348 | 1.694 | 0.6486 | 0.2428 |
| 95% CI of median | | | | | | |
| Actual confidence le | 95.90% | 95.30% | 96.85% | 96.15% | 97.06% | 97.25% |
| Lower confidence lir | 0.05793 | 1.446 | 0.5141 | 0.1368 | 0.04727 | 0.00982 |
| Upper confidence lir | 0.1257 | 2.109 | 1.596 | 0.3618 | 0.1764 | 0.01882 |
| Mean | 0.1244 | 2.541 | 1.808 | 0.3092 | 0.1497 | 0.02403 |
| Std. Deviation | 0.102 | 3.186 | 2.201 | 0.3349 | 0.1412 | 0.03835 |
| Std. Error of Mean | 0.01725 | 0.5237 | 0.3356 | 0.05295 | 0.02535 | 0.00599 |

**Supplementary Table 2:** Statistical significance of the differences in total zinc from Brown-Forsythe and Welch ANOVA with Dunnett's T3 multiple comparison test between MCF10a and different breast cancer cells in different zinc conditions via ICP-MS analysis.

| Dunnett's T3 multiple comparisons test | Mean Diff. | 95.00% CI of diff | Summary | Adjusted P Value |
|---|---|---|---|---|
| MCF10a MM vs. MDA-MB-231 150ZR | 14.74 | -70.68 to 100.2 | ns | 0.937 |
| MCF10a MM vs. T-47D 150ZR | -0.698 | -111.4 to 110.0 | ns | >0.9999 |
| MCF10a MM vs. MCF7 150ZR | -256.5 | -350.6 to -162.4 | *** | 0.0003 |
| MCF10a MM vs. SK-Br-3 150ZR | -457.7 | -1116 to 200.3 | ns | 0.1287 |
| MCF10a MM vs. MDA-MB-157 150ZR | -9.191 | -86.02 to 67.64 | ns | 0.9981 |
| MDA-MB-231 MM vs. MDA-MB-231 150ZR | -22.74 | -43.72 to -1.754 | * | 0.0364 |
| MDA-MB-157 MM vs. MDA-MB-157 150ZR | -13.23 | -49.37 to 22.90 | ns | 0.6148 |
| SK-Br-3 MM vs. SK-Br-3 150ZR | -469.3 | -1122 to 183.9 | ns | 0.1188 |
| MCF7 MM vs. MCF7 150ZR | -284.8 | -376.6 to -193.0 | *** | 0.0003 |
| T-47D MM vs. T-47D 150ZR | -33.03 | -138.9 to 72.89 | ns | 0.8218 |
| MDA-MB-231 MM vs. MDA-MB-157 MM | -33.44 | -62.39 to -4.485 | * | 0.025 |
| MDA-MB-231 MM vs. MCF7 150ZR | -294 | -453.9 to -134.1 | ** | 0.0087 |
| MDA-MB-157 MM vs. MCF7 150ZR | -260.5 | -419.1 to -101.9 | * | 0.0121 |
| SK-Br-3 MM vs. MCF7 150ZR | -268 | -428.4 to -107.6 | * | 0.0115 |
| MCF7 MM vs. MCF7 150ZR | -284.8 | -414.8 to -154.8 | ** | 0.0014 |
| T-47D MM vs. MCF7 150ZR | -288.8 | -424.7 to -152.9 | ** | 0.0016 |
| MDA-MB-231 30ZR vs. MCF7 150ZR | -276.8 | -433.3 to -120.3 | ** | 0.0098 |
| MDA-MB-157 30ZR vs. MCF7 150ZR | -260.2 | -396.7 to -123.8 | ** | 0.0028 |
| SK-Br-3 30ZR vs. MCF7 150ZR | -268.2 | -429.8 to -106.7 | * | 0.0118 |
| MCF7 30ZR vs. MCF7 150ZR | -274.3 | -403.9 to -144.7 | ** | 0.0017 |
| T-47D 30ZR vs. MCF7 150ZR | -282.5 | -413.4 to -151.7 | ** | 0.0015 |
| MDA-MB-231 150ZR vs. MCF7 150ZR | -271.2 | -429.2 to -113.3 | * | 0.0107 |
| MDA-MB-157 150ZR vs. MCF7 150ZR | -247.3 | -383.3 to -111.3 | ** | 0.0055 |
| MCF7 150ZR vs. T-47D 150ZR | 255.8 | 97.34 to 414.2 | ** | 0.0043 |

**Supplementary Table 3:** Total amount of non-zinc trace elements in non-cancerous MCF10A cells and different breast cancer cells in different zinc conditions via ICP-MS analysis.

| Cell types | | Total Cu fg/cell | Total Fe fg/cell | Total Mn fg/cell | Total Ca pg/cell | Total Co ag/cell | Total Mg pg/cell | Total P pg/cell | Total S pg/cell |
|---|---|---|---|---|---|---|---|---|---|
| MCF10a | MM | 4.2 ± 2.0 | 27.1 ± 11.0 | 0.6 ± 0.2 | 0.02 ± 0.0 | 19.7 ± 8.3 | 0.46 ± 0.18 | 10.7 ± 4.6 | 4.4 ± 1.8 |
| | 30ZR | 5.1 ± 4.1 | 22.7 ± 18.0 | 0.5 ± 0.3 | 0.03 ± 0.03 | 16.0 ± 11.6 | 0.46 ± 0.28 | 11.0 ± 8.3 | 4.8 ± 3.4 |
| | 150ZR | n.a. | n.a | n.a. | n.a. | n.a. | n.a. | n.a. | n.a. |
| MDA-MB-231 | MM | 1.43 ± 0.3 | 9.4 ± 1.5 | 2.4 ± 0.5 | 0.04 ± 0.01 | 5.1 ± 2.6 | 0.66 ± 0.15 | 11.4 ± 2.3 | 5.2 ± 1.0 |
| | 30ZR | 1.7 ± 0.3 | 9.0 ± 0.9 | 2.0 ± 0.1 | 0.05 ± 0.01 | 6.0 ± 4.8 | 0.65 ± 0.02 | 11.7 ± 0.7 | 5.3 ± 0.1 |
| | 150ZR | 1.6 ± 0.1 | 10.6 ± 2.0 | 1.4 ± 0.1 | 0.05 ± 0.01 | 3.9 ± 0.1 | 0.66 ± 0.03 | 11.2 ± 1.7 | 5.5 ± 0.8 |
| MDA-MB-157 | MM | 3.8 ± 0.9 | 26.6 ± 4.1 | 4.0 ± 0.6 | 0.27 ± 0.07 | 23.4 ± 4.6 | 1.05 ± 0.21 | 27.2 ± 18.9 | 7.4 ± 0.9 |
| | 30ZR | 2.9 ± 0.9 | 27.0 ± 9.4 | 3.2 ± 1.0 | 0.18 ± 0.05 | 26.2 ± 13.1 | 0.99 ± 0.33 | 29.9 ± 15.4 | 7.0 ± 2.2 |
| | 150ZR | 2.9 ± 0.5 | 28.8 ± 13.0 | 2.3 ± 0.4 | 0.42 ± 0.4 | 26.5 ± 18.0 | 0.99 ± 0.17 | 27.6 ± 8.5 | 6.9 ± 1.3 |
| MCF7 | MM | 2.6 ± 1.2 | 29.1 ± 8.1 | 1.1 ± 0.3 | 0.06 ± 0.01 | 5.8 ± 1.9 | 0.66 ± 0.29 | 13.2 ± 4.2 | 6.6 ± 2.3 |
| | 30ZR | 6.4 ± 7.8 | 34.7 ± 9.0 | 0.7 ± 0.2 | 0.11 ± 0.05 | 8.4 ± 2.9 | 0.72 ± 0.25 | 15.0 ± 5.6 | 7.3 ± 2.8 |
| | 150ZR | 6.1 ± 0.4 | 30.6 ± 4.4 | 0.5 ± 0.1 | 0.07 ± 0.01 | 7.6 ± 2.9 | 0.77 ± 0.15 | 14.1 ± 1.6 | 8.4 ± 1.1 |
| SK-Br-3 | MM | 3.5 ± 0.14 | 21.2 ± 4.9 | 2.4 ± 1.1 | 0.07 ± 0.01 | 42.6 ± 5.3 | 1.00 ± 0.10 | 20.4 ± 4.9 | 7.0 ± 1.5 |
| | 30ZR | 3.3 ± 0.5 | 18.7 ± 3.4 | 1.3 ± 0.3 | 0.06 ± 0.01 | 36.5 ± 5.9 | 0.85 ± 0.14 | 17.1 ± 2.8 | 6.2 ± 1.2 |
| | 150ZR | 4.4 ± 1.1 | 22.8 ± 4.8 | 0.9 ± 0.3 | 0.06 ± 0.01 | 43.7 ± 7.9 | 1.00 ± 0.23 | 20.8 ± 4.2 | 8.2 ± 1.7 |
| T-47D | MM | 6.71 ± 4.2 | 14.9 ± 7.6 | 2.1 ± 1.3 | 0.22 ± 0.14 | 25.4 ± 28.2 | 0.78 ± 0.39 | 8.9 ± 5.9 | 7.6 ± 3.9 |
| | 30ZR | 7.2 ± 3.7 | 13.4 ± 4.4 | 1.0 ± 0.6 | 0.13 ± 0.07 | 15.6 ± 9.2 | 0.53 ± 0.16 | 10.7 ± 3.0 | 5.3 ± 1.4 |
| | 150ZR | 6.9 ± 3.0 | 18.5 ± 10.5 | 0.4 ± 0.1 | 0.66 ± 0.37 | 70.3 ± 40.1 | 0.09 ± 0.06 | 24.5 ± 16.7 | 2.7 ± 0.7 |

| Cell types | Conditions | Total Cu fg/cell | Total Fe fg/cell | Total Mn fg/cell | Total Ca pg/cell | Total Co pg/cell | Total Mg pg/cell | Total P pg/cell | Total S pg/cell |
|---|---|---|---|---|---|---|---|---|---|
| MCF10a | MM | 4.2 ± 2.0 | 27.1 ± 11.0 | 0.6 ± 0.2 | 0.02 ± 0.0 | 19.7 ± 8.3 | 0.46 ± 0.18 | 10.7 ± 4.6 | 4.4 ± 1.8 |
| | 30ZR | 5.1 ± 4.1 | 22.7 ± 18.0 | 0.5 ± 0.3 | 0.03 ± 0.03 | 16.0 ± 11.6 | 0.46 ± 0.28 | 11.0 ± 8.3 | 4.8 ± 3.4 |
| | 150ZR | n.a. | n.a | n.a. | n.a. | n.a. | n.a. | n.a. | n.a. |
| MDA-MB-231 | MM | 1.43 ± 0.3 | 9.4 ± 1.5 | 2.4 ± 0.5 | 0.04 ± 0.01 | 5.1 ± 2.6 | 0.66 ± 0.15 | 11.4 ± 2.3 | 5.2 ± 1.0 |
| | 30ZR | 1.7 ± 0.3 | 9.0 ± 0.9 | 2.0 ± 0.1 | 0.05 ± 0.01 | 6.0 ± 4.8 | 0.65 ± 0.02 | 11.7 ± 0.7 | 5.3 ± 0.1 |
| | 150ZR | 1.6 ± 0.1 | 10.6 ± 2.0 | 1.4 ± 0.1 | 0.05 ± 0.01 | 3.9 ± 0.1 | 0.66 ± 0.03 | 11.2 ± 1.7 | 5.5 ± 0.8 |
| MDA-MB-157 | MM | 3.8 ± 0.9 | 26.6 ± 4.1 | 4.0 ± 0.6 | 0.27 ± 0.07 | 23.4 ± 4.6 | 1.05 ± 0.21 | 27.2 ± 18.9 | 7.4 ± 0.9 |
| | 30ZR | 2.9 ± 0.9 | 27.0 ± 9.4 | 3.2 ± 1.0 | 0.18 ± 0.05 | 26.2 ± 13.1 | 0.99 ± 0.33 | 29.9 ± 15.4 | 7.0 ± 2.2 |
| | 150ZR | 2.9 ± 0.5 | 28.8 ± 13.0 | 2.3 ± 0.4 | 0.42 ± 0.4 | 26.5 ± 18.0 | 0.99 ± 0.17 | 27.6 ± 8.5 | 6.9 ± 1.3 |
| MCF7 | MM | 2.6 ± 1.2 | 29.1 ± 8.1 | 1.1 ± 0.3 | 0.06 ± 0.01 | 5.8 ± 1.9 | 0.66 ± 0.29 | 13.2 ± 4.2 | 6.6 ± 2.3 |
| | 30ZR | 6.4 ± 7.8 | 34.7 ± 9.0 | 0.7 ± 0.2 | 0.11 ± 0.05 | 8.4 ± 2.9 | 0.72 ± 0.25 | 15.0 ± 5.6 | 7.3 ± 2.8 |
| | 150ZR | 6.1 ± 0.4 | 30.6 ± 4.4 | 0.5 ± 0.1 | 0.07 ± 0.01 | 7.6 ± 2.9 | 0.77 ± 0.15 | 14.1 ± 1.6 | 8.4 ± 1.1 |
| SK-Br-3 | MM | 3.5 ± 0.14 | 21.2 ± 4.9 | 2.4 ± 1.1 | 0.07 ± 0.01 | 42.6 ± 5.3 | 1.00 ± 0.10 | 20.4 ± 4.9 | 7.0 ± 1.5 |
| | 30ZR | 3.3 ± 0.5 | 18.7 ± 3.4 | 1.3 ± 0.3 | 0.06 ± 0.01 | 36.5 ± 5.9 | 0.85 ± 0.14 | 17.1 ± 2.8 | 6.2 ± 1.2 |
| | 150ZR | 4.4 ± 1.1 | 22.8 ± 4.8 | 0.9 ± 0.3 | 0.06 ± 0.01 | 43.7 ± 7.9 | 1.00 ± 0.23 | 20.8 ± 4.2 | 8.2 ± 1.7 |
| T-47D | MM | 6.71 ± 4.2 | 14.9 ± 7.6 | 2.1 ± 1.3 | 0.22 ± 0.14 | 25.4 ± 28.2 | 0.78 ± 0.39 | 8.9 ± 5.9 | 7.6 ± 3.9 |
| | 30ZR | 7.2 ± 3.7 | 13.4 ± 4.4 | 1.0 ± 0.6 | 0.13 ± 0.07 | 15.6 ± 9.2 | 0.53 ± 0.16 | 10.7 ± 3.0 | 5.3 ± 1.4 |
| | 150ZR | 6.9 ± 3.0 | 18.5 ± 10.5 | 0.4 ± 0.1 | 0.66 ± 0.37 | 70.3 ± 40.1 | 0.09 ± 0.06 | 24.5 ± 16.7 | 2.7 ± 0.7 |

**Supplementary Table 4:** Statistical significance of the differences in total non-zinc trace elements between MCF10a and different breast cancer cells in different zinc conditions via ICP-MS analysis.

| **Brown-Forsythe and Welch ANOVA with Dunnett's T3 multiple comparisons test** | **Mean Diff.** | **95.00% CI of diff.** | **Summary** | **Adjusted P Value** |
|---|---|---|---|---|
| **Cu** | | | | |
| SK-Br-3 MM vs. MDA-MB-231-30ZR | 1.84 | 0.5504 to 3.130 | * | 0.0138 |
| SK-Br-3 MM vs. MDA-MB-231-150 ZR | 1.92 | 1.452 to 2.396 | **** | <0.0001 |
| SK-Br-3 MM vs. MCF7 150ZR | -2.61 | -3.981 to -1.235 | ** | 0.0046 |
| MDA-MB-231 30ZR vs. MCF7 150ZR | -4.45 | -5.885 to -3.011 | *** | 0.0001 |
| SK-Br-3 MM vs. MCF7 150ZR | -2.81 | -4.864 to -0.752 | * | 0.013 |
| MDA-MB-231 150 ZR vs. MCF7 150ZR | -4.53 | -6.142 to -2.923 | ** | 0.0025 |
| MDA-MB-157 150 ZR vs. MCF7 150ZR | -3.19 | -4.920 to -1.460 | ** | 0.0021 |
| **Fe** | | | | |
| MDA-MB-231 MM vs. MDA-MB-157 MM | -17.2 | -32.22 to -2.258 | * | 0.0304 |
| MDA-MB-231 MM vs. MCF7 150 ZR | -21.27 | -37.31 to -5.243 | * | 0.018 |
| MDA-MB-157 MM vs. MDA-MB-231 30ZR | 17.62 | 0.01445 to 35.22 | * | 0.0499 |
| MDA-MB-157 MM vs. MDA-MB-231-150 ZR | 15.98 | 0.2680 to 31.69 | * | 0.0471 |
| MDA-MB-231-30ZR vs. MCF7 150 ZR | -21.7 | -40.59 to -2.715 | * | 0.0344 |
| MDA-MB-231-150 ZR vs. MCF7 150 ZR | -20.0 | -36.74 to -3.299 | * | 0.0263 |
| **Mn** | | | | |
| MCF10a MM vs. MDA-MB-231 MM | -1.84 | -3.554 to -0.1177 | * | 0.0396 |
| MCF10a MM vs. MDA-MB-157 MM | -3.43 | -5.621 to -1.241 | ** | 0.0097 |
| MCF10a MM vs. MDA-MB-231 30ZR | -1.38 | -2.054 to -0.7032 | ** | 0.0035 |
| MCF10a MM vs. MDA-MB-231 150 ZR | -0.78 | -1.483 to -0.08513 | * | 0.0333 |
| MCF10a MM vs. MDA-MB-157 150 ZR | -1.76 | -3.287 to -0.2294 | * | 0.0304 |
| MDA-MB-231 MM vs. MCF10a 30ZR | 1.931 | 0.3152 to 3.546 | * | 0.021 |
| MDA-MB-231 MM vs. MCF7 30ZR | 1.679 | 0.08132 to 3.277 | * | 0.0407 |
| MDA-MB-231 MM vs. T-47D 150ZR | 2.041 | 0.08155 to 4.000 | * | 0.0446 |
| MDA-MB-157 MM vs. MCF7 MM | 2.864 | 0.7481 to 4.979 | * | 0.0135 |
| MDA-MB-157 MM vs. MCF10a 30ZR | 3.526 | 1.400 to 5.653 | ** | 0.0053 |
| MDA-MB-157 MM vs. MCF10a 30ZR | 3.526 | 1.400 to 5.653 | ** | 0.0053 |
| MDA-MB-157 MM vs. SK-Br-3 30ZR | 2.707 | 0.6104 to 4.803 | * | 0.0167 |
| MDA-MB-157 MM vs. MCF7 30ZR | 3.274 | 1.016 to 5.533 | * | 0.013 |
| MDA-MB-157 MM vs. T-47D 30ZR | 3.032 | 0.6651 to 5.399 | * | 0.0148 |
| MDA-MB-157 MM vs. MDA-MB-231-150ZR | 2.647 | 0.06801 to 5.226 | * | 0.0465 |
| MDA-MB-157 MM vs. SK-Br-3 150ZR | 3.084 | 0.9894 to 5.178 | ** | 0.0092 |
| MDA-MB-157 MM vs. MCF7 150 ZR | 3.461 | 0.8833 to 6.040 | * | 0.0217 |
| MDA-MB-157 MM vs. T-47D 150ZR | 3.636 | 1.067 to 6.205 | * | 0.0187 |
| MCF10a 30ZR vs. MDA-MB-231-30ZR | -1.47 | -2.94 to -0.01 | * | 0.0491 |
| MCF10a 30ZR vs. MDA-MB-157 150 ZR | -1.85 | -3.34 to -0.37 | * | 0.0168 |

| | | | | |
|---|---|---|---|---|
| MDA-MB-231-30ZR vs. MCF7 30ZR | 1.222 | 0.1537 to 2.290 | * | 0.0343 |
| MDA-MB-231-30ZR vs. MDA-MB-231-150ZR | 0.595 | 0.3105 to 0.8788 | ** | 0.0018 |
| MDA-MB-231-30ZR vs. MCF7 150 ZR | 1.409 | 1.151 to 1.668 | **** | <0.0001 |
| MDA-MB-231-30ZR vs. T-47D 150ZR | 1.584 | 1.371 to 1.797 | **** | <0.0001 |
| MCF7 30ZR vs. MDA-MB-157 150 ZR | -1.60 | -3.041 to -0.1623 | * | 0.0319 |
| MDA-MB-231-150 ZR vs. MCF7 150 ZR | 0.815 | 0.5174 to 1.112 | *** | 0.0002 |
| MDA-MB-231-150 ZR vs. T-47D 150ZR | 0.989 | 0.7087 to 1.270 | *** | 0.0001 |
| MDA-MB-157 150 ZR vs. MCF7 150ZR | 1.789 | 0.06613 to 3.511 | * | 0.045 |
| MDA-MB-157 150 ZR vs. T-47D 150ZR | 1.963 | 0.2548 to 3.671 | * | 0.0339 |
| **Ca** | | | | |
| MCF10a MM vs. SK-Br-3 MM | -0.05 | -0.0777 to -0.021 | ** | 0.0043 |
| MCF10a MM vs. MDA-MB-231-30ZR | -0.03 | -0.056 to -0.0004 | * | 0.0473 |
| MCF10a MM vs. MDA-MB-231-150 ZR | -0.03 | -0.051 to -0.0071 | * | 0.0124 |
| **Co** | | | | |
| MDA-MB-231 MM vs. MDA-MB-157 MM | -0.018 | -0.0346 to -0.002 | * | 0.03 |
| MDA-MB-231 MM vs. SK-Br-3 MM | -0.038 | -0.0577 to -0.017 | ** | 0.0051 |
| MDA-MB-231 MM vs. SK-Br-3 30ZR | -0.031 | -0.0537 to -0.009 | * | 0.0145 |
| MDA-MB-231 MM vs. SK-Br-3 150ZR | -0.039 | -0.0673 to -0.001 | * | 0.0173 |
| MDA-MB-157 MM vs. MCF7 MM | 0.018 | 0.0003 to 0.035 | * | 0.0471 |
| SK-Br-3 MM vs. MCF7 MM | 0.037 | 0.01752 to 0.056 | ** | 0.0046 |
| SK-Br-3 MM vs. MDA-MB-231 30ZR | 0.037 | 0.01652 to 0.057 | ** | 0.0022 |
| SK-Br-3 MM vs. MCF7 30ZR | 0.034 | 0.01580 to 0.053 | ** | 0.0031 |
| SK-Br-3 MM vs. MDA-MB-231-150 ZR | 0.039 | 0.01620 to 0.061 | * | 0.0106 |
| SK-Br-3 MM vs. MCF7 150 ZR | 0.035 | 0.01658 to 0.054 | ** | 0.0028 |
| MCF7 MM vs. SK-Br-3 30ZR | -0.030 | -0.0522 to -0.00919 | * | 0.0137 |
| MCF7 MM vs. SK-Br-3 150ZR | -0.037 | -0.0723 to -0.00350 | * | 0.0381 |
| MDA-MB-231-30ZR vs. SK-Br-3 30ZR | -0.030 | -0.0520 to -0.0089 | ** | 0.0087 |
| MDA-MB-231-30ZR vs. SK-Br-3 150ZR | -0.038 | -0.066 to -0.009 | * | 0.0146 |
| SK-Br-3 30ZR vs. MCF7 30ZR | 0.028 | 0.0052 to 0.051 | * | 0.0239 |
| SK-Br-3 30ZR vs. MDA-MB-231 150 ZR | 0.033 | 0.007 to 0.058 | * | 0.0245 |
| SK-Br-3 30ZR vs. MCF7 150 ZR | 0.029 | 0.00603 to 0.0518 | * | 0.0216 |
| MCF7 30ZR vs. SK-Br-3 150ZR | -0.035 | -0.064 to -0.006 | * | 0.0253 |
| MDA-MB-231-150ZR vs. SK-Br-3 150ZR | -0.040 | -0.073 to -0.006 | * | 0.0312 |
| SK-Br-3 150ZR vs. MCF7 150 ZR | 0.036 | 0.0069 to 0.065 | * | 0.0233 |
| **P** | | | | |
| no significance | | | | |
| **S** | | | | |

| | | | | |
|---|---|---|---|---|
| MDA-MB-157 MM vs. T-47D 150ZR | 4.675 | 1.377 to 7.973 | ** | 0.0087 |
| MCF7 150 ZR vs. T-47D 150ZR | 5.698 | 1.492 to 9.903 | * | 0.0135 |
| **Mg** | | | | |
| MDA-MB-157 MM vs. T-47D 150ZR | 0.955 | 0.190 to 1.720 | * | 0.0225 |
| SK-Br-3 MM vs. T-47D 150ZR | 0.907 | 0.519 to 1.294 | ** | 0.001 |
| MDA-MB-231-30ZR vs. T-47D 150ZR | 0.552 | 0.097 to 1.007 | * | 0.0342 |
| SK-Br-3 30ZR vs. T-47D 150ZR | 0.755 | 0.197 to 1.313 | * | 0.0167 |
| MDA-MB-231-150ZR vs. T-47D 150ZR | 0.563 | 0.251 to 0.874 | ** | 0.0092 |
| MDA-MB-157 150ZR vs. T-47D 150ZR | 0.893 | 0.260 to 1.525 | * | 0.0143 |
| MCF7 150 ZR vs. T-47D 150ZR | 0.67 | 0.103 to 1.237 | * | 0.0276 |
